# Systems Level Analysis of Gene, Pathway and Phytochemical Associations with Psoriasis

**DOI:** 10.64898/2025.12.20.695744

**Authors:** Sonalika Ray, Ojasvi Dutta, Neoklis Apostolopoulos, Konstantin G. Kousoulas, Jean Christopher Chamcheu, Rishemjit Kaur

**Affiliations:** CSIR-Central Scientific Instruments organization, Chandigarh, India; Academy of Scientific and Innovative Research (AcSIR), Ghaziabad, India; Department of Pathobiological Sciences, School of Veterinary Medicine, Louisiana State University, Baton Rouge, Louisiana-70803; Division of Biotechnology and Molecular Medicine, School of Veterinary Medicine, Louisiana State University, Baton Rouge, Louisiana-70803; Department of Veterinary Clinical Sciences, School of Veterinary Medicine, Louisiana State University.

**Keywords:** Psoriasis, Transcriptomics, Interferon signaling, Upstream regulators, Immunometabolic pathways, Network pharmacology, Phytochemicals, Synergy analysis, Multi-target therapy

## Abstract

Psoriasis is an inflammatory skin disorder driven by abnormal immune activation that promotes excessive proliferation and accelerated turnover of epidermal keratinocytes. IL-17 and TNF pathways are well established in psoriasis, but the other mechanisms that keep the disease active and link it to systemic comorbidities are not yet fully understood. A combined transcriptomic and systems biology framework was applied to map regulatory circuits in psoriatic lesions and to identify phytochemical candidates capable of multi-target modulation for topical intervention. Differential gene expression between lesional and healthy skin was analyzed, followed by functional characterization, employing Qiagen’s Ingenuity Pathway Analysis (IPA) for pathway and upstream regulator inference, protein-protein interaction network, and chemical-gene interaction mapping. This integrative strategy revealed a transcriptional landscape dominated by type I/III interferon signaling, antiviral and antimicrobial responses, immunometabolic dysregulation, and transcriptional hubs centered on AP-1 and CREB1. Several previously unreported genes and upstream regulators without prior documented association with psoriasis were identified within inflammatory and cell migration-related modules, indicating unexplored regulatory layers in disease control. Network-guided chemical prioritization and direction-of-effect filtering highlighted seven phytochemicals (mahanine, atractylon, protopine, annomontine, taraxasterol, tricin, and tamarixetin) with multi-target activity across key disease axes. ADMET-based screening suggested protopine and atractylon as favorable candidates for topical delivery, while synergy modeling identified compatible phytochemical combinations, with flavonoid-alkaloid pairings among the top candidates. This multi-layered approach provides mechanistically informed phytochemicals targeting the IL-17/TNF-interferon-AP-1/CREB1-COX-2/MMP9 axis in psoriasis. Experimental validation in keratinocyte and organotypic skin models will be required to determine whether these compounds, individually or in combination, can effectively modulate psoriatic signaling in vivo.

## 1 Introduction

Plants have served as a primary source of biologically active compounds throughout human history, and their chemical diversity constitutes one of the largest naturally occurring reservoirs of molecules with therapeutic relevance [1]. The secondary metabolites produced by plants collectively termed phytochemicals, span structurally distinct classes including flavonoids, terpenoids, alkaloids, coumarins, and polyphenols, each with characteristic mechanisms of cellular and molecular action [2]. What differentiates phytochemicals from narrowly targeted agents is their inherent capacity to modulate multiple biological pathways simultaneously, a property that arises from the structural complexity and functional versatility of plant-derived molecules [1]. This multi-target behavior is particularly relevant for diseases governed by interconnected regulatory networks, where perturbation of a single node is often insufficient to achieve durable biological effects [3]. Traditional medical systems such as Ayurveda, Siddha, and Traditional Chinese Medicine have exploited this property for centuries, using complex polyherbal preparations to manage chronic inflammatory conditions with a favorable long-term safety record [4, 5].

Chronic inflammatory skin diseases represents a particularly active area of phytochemical research, given that skin is both accessible for topical delivery and subject to complex immunological dysregulation that benefits from multi-target modulation [2]. Phytochemicals have been documented to regulate inflammatory signaling, support epidermal barrier integrity, attenuate oxidative stress, and modulate cutaneous immune responses across a range of skin conditions [1]. Several specific compounds have been characterized in detail: curcumin from *Curcuma longa* [4], aloe emodin from *Aloe vera* [4], and psoralen from *Psoralea corylifolia* [6] each show well-documented anti-inflammatory and anti-proliferative effects relevant to chronic epidermal pathology. Plant sources including *Nigella sativa*, *Rubia cordifolia*, *Smilax china*, *Thespesia populnea*, and *Wrightia tinctoria* [7, 8] have similarly demonstrated immunomodulatory properties associated with improved outcomes in inflammatory skin disorders. Flavonoids and polyphenolic compounds, in particular, suppress NF-*κ*B activation, reduce pro-inflammatory cytokine output, and inhibit abnormal keratinocyte proliferation - processes central to chronic skin inflammation [2]. Despite this breadth of documented activity, fewer than 15% of plant-derived molecules have been formally evaluated for dermatological applications, reflecting the absence of systematic frameworks to link phytochemical activity to specific disease gene networks [1].

Modern computational approaches have begun to close this gap. Network pharmacology, transcriptomics-guided target identification, molecular docking, and ADMET profiling together provide a structured route from disease gene signatures to rationally prioritized phytochemical candidates [3, 9]. Applied to psoriasis, this strategy has already produced mechanistically grounded findings: quercetin was shown through network analysis to simultaneously modulate Notch1, PI3K/Akt, and Glut-1 signaling in keratinocytes [10]; a synergistic interaction between baicalin and echinacoside was computationally predicted and experimentally confirmed to suppress IL-17 and TNF signaling more effectively in combination than as individual agents [11]; and oleanolic acid was validated against psoriasis-relevant targets including MAPK3, STAT3, PPARG, and PTGS2 in an imiquimod model [12]. These examples demonstrate that integrating transcriptomic data with phytochemical interaction databases can generate testable, mechanistically supported candidate hypotheses for complex inflammatory skin diseases.

Among such diseases, psoriasis is one of the most molecularly well-characterized and clinically significant targets for phytochemical investigation. It is a chronic autoimmune skin disorder affecting approximately 2-3% of the global population, with higher prevalence among people of European descent than among Asian or African populations [13, 14], and with disease onset concentrated in two epidemiological peaks: early adulthood (20-30 years) and later adulthood (50-60 years) [14]. Its pathogenesis involves a continuous interaction between genetic susceptibility and environmental triggers including infection, physical skin trauma, psychological stress, and certain external agents [15, 16]. At the cellular level, dendritic cells activate Th1 and Th17 lymphocytes together with plasmacytoid cells that produce type I interferons; these populations collectively release TNF-*α*, IL-17, IL-23, and interferon-*γ*, driving keratinocyte hyperproliferation and sustaining a self-reinforcing inflammatory cycle [17, 18].Beyond the cytokine axes that dominate current literature, the molecular landscape of psoriasis also encompasses antiviral and type I/III interferon programs, cell cycle dysregulation, oxidative stress, and immunometabolic disruption [19, 16]. Inflammatory mediators released from lesional skin enter systemic circulation and contribute to comorbidities including cardiovascular disease, metabolic syndrome, and inflammatory bowel disease [20, 21], reducing quality of life to levels comparable to those associated with other major chronic conditions [22]. This multi-module regulatory organization makes psoriasis a well-suited disease context for phytochemicals capable of engaging several pathological nodes concurrently.

From a delivery perspective, phytochemicals also offer practical advantages for skin-directed intervention. Their physicochemical diversity permits formulation into nanoparticles, lipid-based carriers, and hydrogels that sustain local activity while limiting systemic exposure - properties important for conditions requiring long-term topical management [2]. Polyherbal preparations such as *Parangichakkai Chooranam*, when analyzed through systems pharmacology frameworks, provide additional precedent for integrative phytochemical analysis in psoriasis [5]. Individual plant-derived compounds used as adjunct agents have similarly shown favorable tolerability profiles in chronic dermatosis settings [23]. Nevertheless, the systematic isolation and functional characterization of phytochemicals specifically mapped to psoriatic gene pathways remains limited, particularly with respect to candidates suited for topical development [3, 24].

The present study integrates transcriptomic analysis with network pharmacology to identify and prioritize phytochemicals targeting regulatory genes in psoriatic lesions, with emphasis on candidates suitable for topical application. Specifically, the study maps DEGs from lesional skin to curated chemical-gene interaction databases, evaluates direction-of-effect consistency across psoriasis-relevant pathways, and assesses ADMET-based topical suitability and pairwise synergy among short-listed candidates. This framework provides a biologically grounded route from disease transcriptomics to mechanism-guided phytochemical prioritization for experimental follow-up in skin model systems.

## 2 Results

We analyzed gene expression profiles from NCBI GEO [25] dataset GSE14905 to define psoriasis-associated transcriptomic signature and prioritize multi-target phytochemicals. The analysis followed a structured workflow (as shown in Methods Section 3), including PCA-based sample quality assessment, differential gene expression (DGE) analysis, evidence-based classification of previously unreported and high-evidence genes, IPA pathway and regulator analysis, chemical-gene interaction mapping, direction-of-effect-based phytochemical prioritization, ADMET screening for topical suitability, in-silico synergy assessment, protein-protein interaction (PPI) network analysis, and construction of a phytochemical-gene-pathway tripartite network. The detailed results from each stage are presented below.

### 2.1 Identification of sample outliers and clusters through PCA

Principal Component Analysis (PCA) was performed on gene expression data from 21 healthy controls and 28 psoriasis samples (Refer to Methods Section 3.1) and showed a clear separation between psoriasis and control groups, Figure 1A. Lesional samples (red) formed a different group from normal samples (green) and non-involved samples (blue), reflecting the transcriptomic differences associated with disease status. Uninvolved skin samples largely overlapped with the healthy controls, indicating minimal transcriptomic alteration in these regions. Three lesional samples deviated from the main lesional group and were identified as outliers. These samples were excluded from further analysis to reduce the variability in downstream results.

**Fig. 1:**
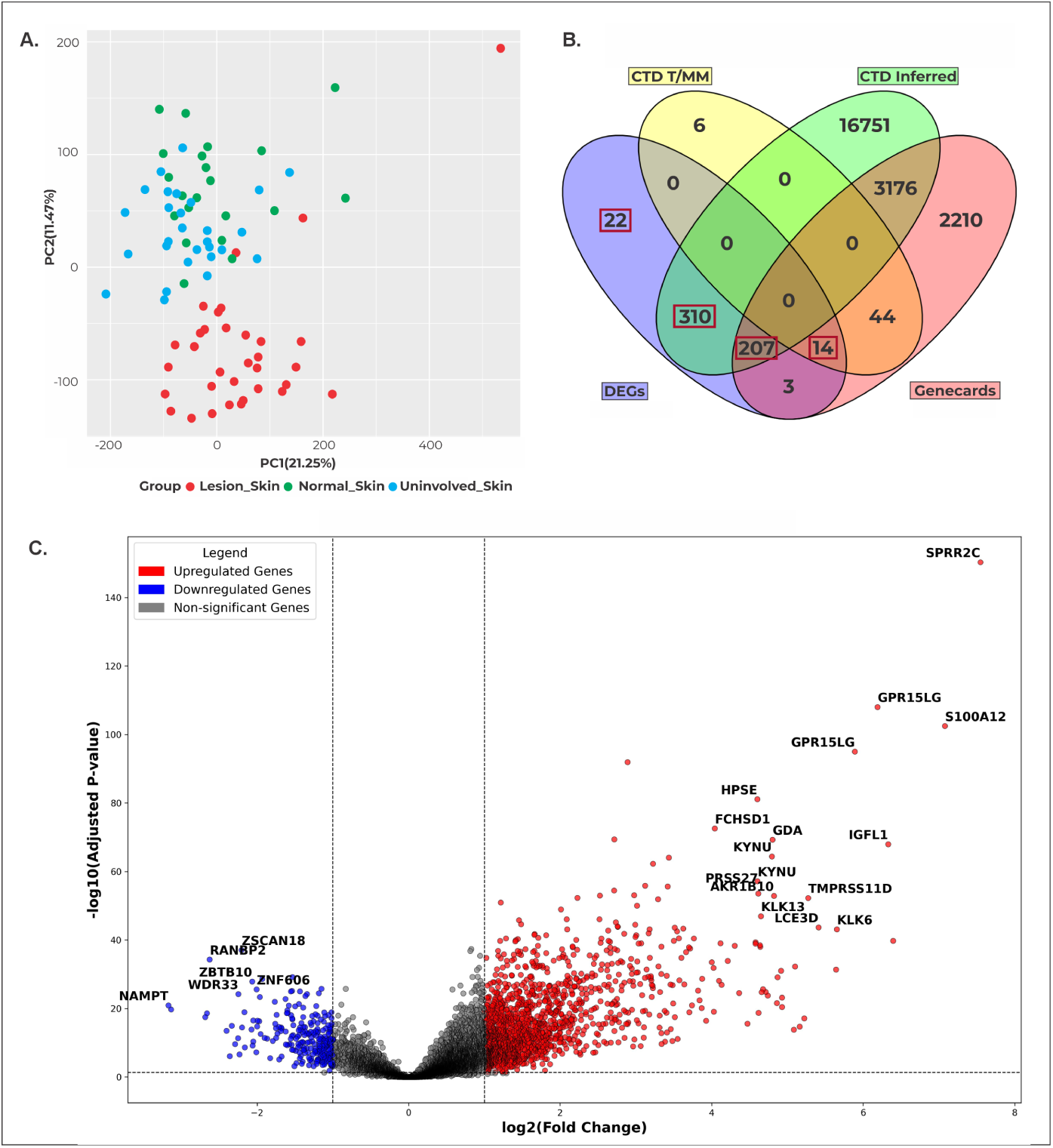
**(A)** PCA plot showing separation of lesional, uninvolved, and healthy skin samples. **(B)** Four-set Venn diagram showing overlap between DEGs identified in this study and psoriasis-associated genes from CTD (Inferred and T/MM) and GeneCards. **(C)** Volcano plot of gene expression differences between lesional and normal skin.

### 2.2 Identification of DEGs

We identified 556 DEGs by comparing lesional and healthy skin samples, which included 270 upregulated and 286 downregulated transcripts (Refer to Methods Section 3.2). The complete DEGs list is provided in Supplementary File 1. The volcano plot (Figure 1C) shows the distribution of DEGs by fold change and statistical significance, with distinct separation of upregulated (red) and downregulated (blue) genes and non-significant genes clustered around the center. The top DEGs with the highest magnitude of fold changes are highlighted. The top upregulated genes included *SPRR2C* (7.549), *S100A12* (7.079), *TCN1* (6.396), *IGFL1* (6.332), *GPR15LG* (6.191), *KLK6* (5.654), *CXCL9* (5.643), *LCE3D* (5.413), *TMPRSS11D* (5.275), and *S100A7A* (5.223), while the top downregulated genes included *PPP1R1A* (−3.770), *STC1* (−3.796), *LPL* (−3.814), *KRT77* (−3.871), *ACVR1C* (−3.952), *IL6* (−4.328), *MUC7* (−4.423), *FOSB* (−4.455), *CYP1A1* (−4.622), and *FOS* (−4.924), indicating clear differences in gene expression between lesional and healthy skin samples.

### 2.3 Identification of unreported and high evidence DEGs

We compared 556 DEGs with psoriasis-associated genes listed in the Comparative Toxicogenomics Database (CTD) [26] and GeneCards [27] (Refer to Methods Section 3.3). CTD contained 20,555 psoriasis-associated genes, including 63 with marker or mechanistic (MM) evidence, 1 with therapeutic (T) evidence and 20,491 inferred associations. GeneCards listed 5,654 psoriasis-related genes. Overlap analysis using Venny [28] identified shared and unique gene sets, as shown in Figure 1B. Three DEG sets were defined (Figure 2):

- Set 1: 14 high-evidence DEGs, defined as genes overlapping both CTD T/MM evidence and GeneCards.
- Set 2: 517 DEGs (310+207) were also found in CTD inferred genes set. Among them, 32 DEGs lacked direct evidence in PubMed or Open Targets, indicating genes with indirect or computationally inferred links to psoriasis. Further we performed functional enrichment analysis and identified 14 DEGs with significant enrichment, with detailed results provided in Appendix A.
- Set 3: 22 exclusive DEGs which have not been reported in CTD (neither marker/mechanistic nor inferred evidence) or GeneCards. These 22 DEGs were further screened using PubMed and the Open Targets platform to verify the presence of any previously reported psoriasis associations. We found 4 previously unreported DEGs, representing genes with no documented association in CTD, GeneCards, PubMed, or Open Targets;

**Fig. 2:**
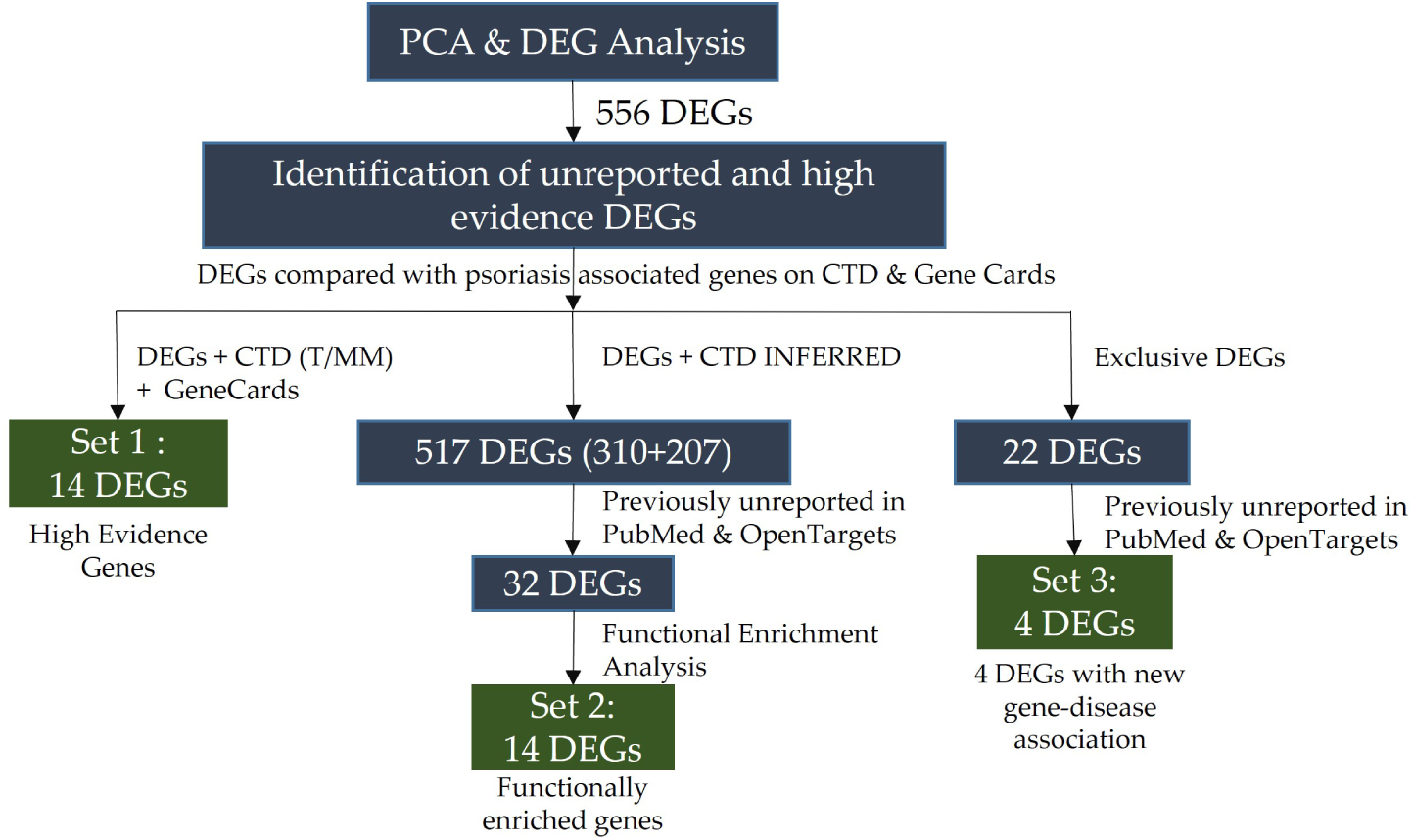
DEG classification into three evidence-stratified sets. Overlap of 556 DEGs with CTD, GeneCards, PubMed, and Open Targets defines Set 1 (14 high-evidence), Set 2 (14 functionally enriched), and Set 3 (4 previously unreported DEGs).

Detailed results are in Supplementary File 1.

### 2.4 Identification of regulatory networks and upstream drivers

We performed functional enrichment analysis on Qiagen’s Ingenuity Pathway Analysis (IPA) on 556 DEGs, with their respective log_2_ fold-change and *p*-values (Refer to Methods Section 3.4) and obtained IPA graphical summary of major regulators and functional interactions, Upstream Regulators (URs), Canonical Pathways, Diseases & Biofunctions, and Machine Learning (ML) Disease Pathways.

#### Upstream regulator analysis

A total of 230 URs satisfied the filtering criteria (*p <* 0.05; activation *z*-score *>* +2 or *<* −2) (see Supplementary File 3). They were used alongside the DEGs to guide phytochemical prioritization. Further, to determine URs with previous unknown association, these were cross-checked against CTD, GeneCards, and PubMed. We found nine such URs: *IFNA6*, *IFNA7*, *IFNA10*, *IFNA14*, *IFNA16*, *IFNA21*, *IFNL1*, *IFNL4*, and *MRGPRA2A*. These spanned two biologically distinct interferon subtypes: (i) type I interferons (*IFNA6*, *IFNA7*, *IFNA10*, *IFNA14*, *IFNA16*, *IFNA21*) and (ii) type III interferons (*IFNL1*, *IFNL4*), alongside *MRGPRA2A*.

#### IPA Graphical summary analysis

The IPA Graphical Summary (Figure 3) integrates key findings from upstream regulator, pathway, and diseases and biofunction analyses into a single consolidated network, highlighting the most statistically significant DEGs and their predicted activation (orange) or inhibition (blue) states. The results showed activation of key interferon-related regulators, including *STAT1*, *IRF1*, *IRF3*, and *IRF7*. *cGAS* and *MAVS* were connected to these regulators, and *IFNA* family members along with *IFNL1* were associated with interferon signaling, antiviral, and antimicrobial response pathways. The only molecule predicted to be inhibited was *F3* (tissue factor/coagulation factor III).

**Fig. 3:**
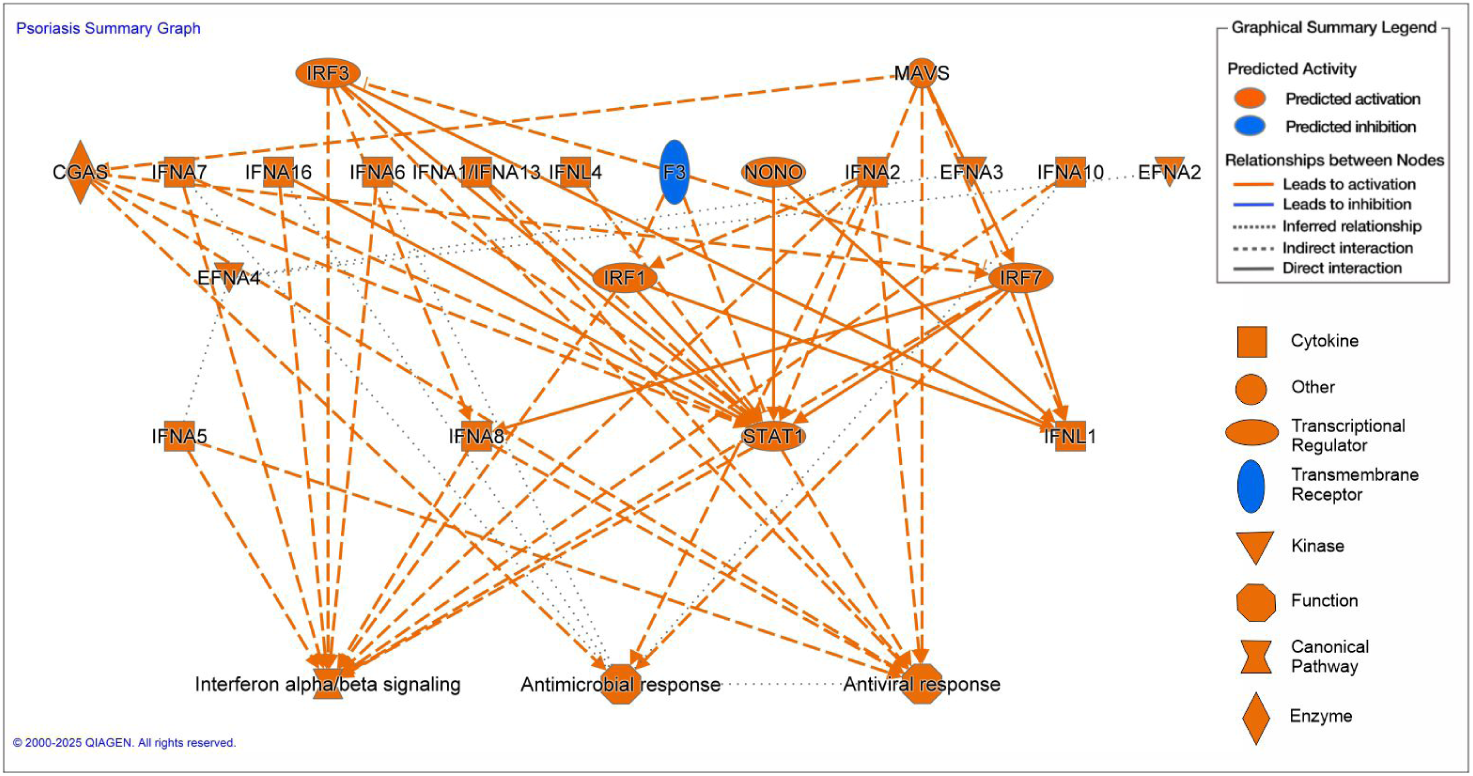
Graphical summary generated by IPA showing predicted upstream regulators and functional interactions in psoriasis.

### 2.5 Identification and prioritization of phytochemicals targeting previously unreported and high-evidence genes

We identified 13 shortlisted phytochemicals targeting Set 1-high evidence psoriasis-associated genes (Refer to Methods Section 3.5) which are annomontine, olivetol, protopine, hypaconitine, taraxasterol, atractylon, carboxyatractyloside, mahanine, bilobalide, robinetin, scutellarein, tamarixetin, and tricin. For Set 2 and Set 3 genes, we found phytochemicals already known for psoriasis (for details, refer to Supplementary File 4). We further checked the direction of effect of the 13 shortlisted phytochemicals. The ones predicted to counteract disease-associated expression patterns were prioritized. This included suppression of overexpressed genes and restoration of under-expressed genes. Based on this screening, 7 phytochemicals were prioritized: mahanine, taraxasterol, tricin, tamarixetin, annomontine, protopine, and atractylon. The gene targets, corresponding fold changes, and reported molecular effects are summarized in Table 1. The phytochemicals showed directionally consistent interactions with key psoriasis-associated genes, including: (i) downregulation of overexpressed proliferative genes (*CCNB1*, *CDK1*); (ii) inhibition of inflammatory and matrix-remodeling mediators (*MMP9*, *PTGS2*);(iii) restoration of detoxification and regulatory genes (*CYP1A1*, *CREB1*). In addition, *CREB1* was identified as an upstream regulator targeting multiple DEGs, including *ADM, APOLD1, ARG1, ATF3, AURKA, BHLHE40, BIRC5, CCN1, CCNA2, CCNB1, CCNB2, CD3D, CHRM3, CXCL8, DUSP1, EGR1, ESRRG, FASN, FLT1, FOS, FOSB, GADD45B, GAL, GDA, GLDC, HSD11B1, IL6, INA, JUN, LCN2, LPL, MKI67, MMP1, NFIL3, NR4A1, NR4A2, PCK1, PER1, PPARG, PPARGC1A, PPP1R15A, PTGS2, RBP4, SIK1, SLC2A3, SMPD3, SOCS2, SOD2, STC1, TF, VEGFA, ZFP36*.

**Table 1:**
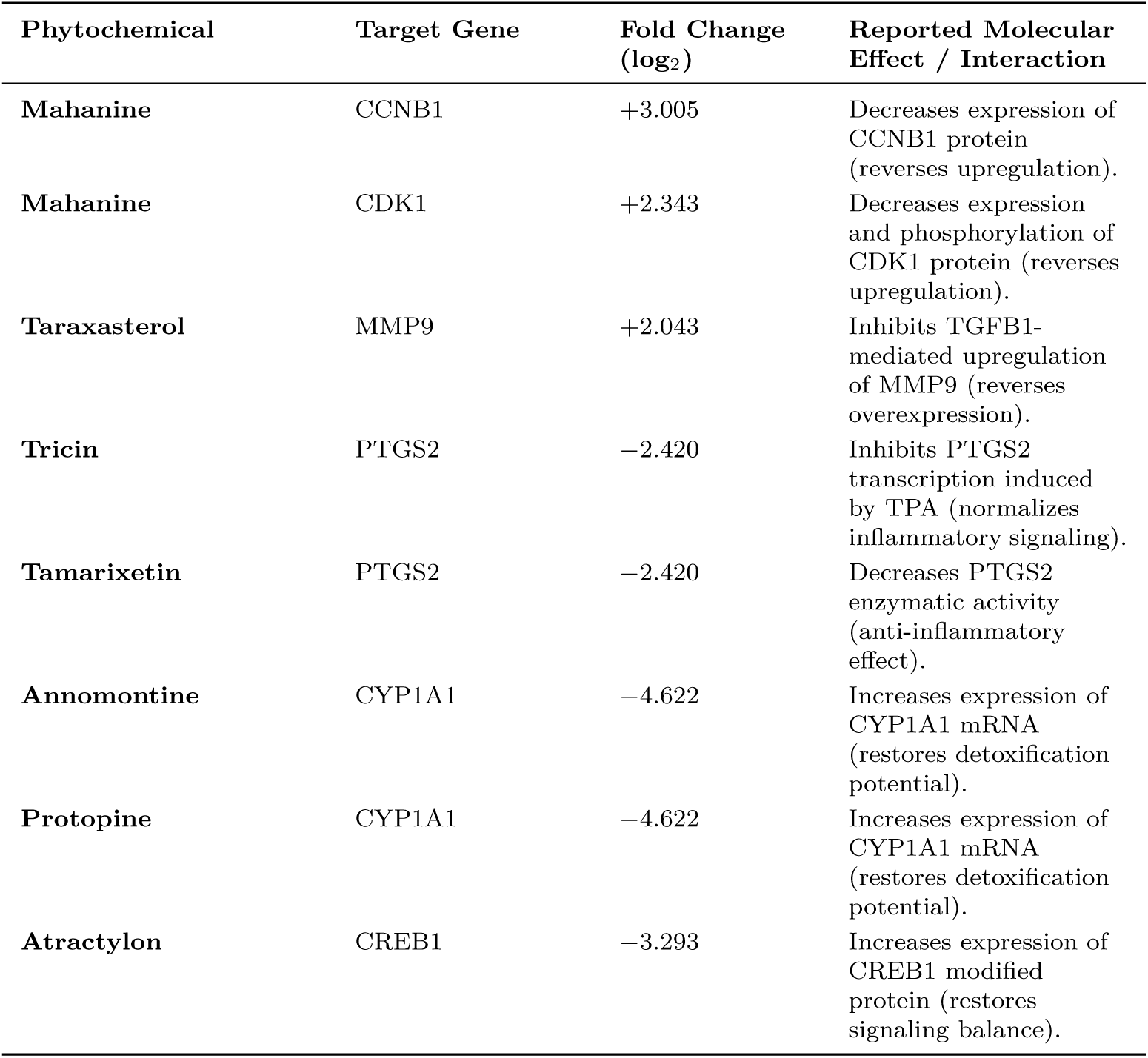
Shortlisted phytochemicals, their target genes, fold-change values, and reported molecular effects. These compounds were prioritized based on direction-of-effect consistency.

#### Evaluation of phytochemicals for topical suitability using ADMET-AI

The 7 prioritized phytochemicals were evaluated for topical suitability using the ADMET-AI platform (Refer to Methods Section 3.6), and all seven were retained as candidates with profiles supporting further development (see Supplementary File 5). Protopine and atractylon demonstrated the most favorable profiles for passive dermal penetration, with moderate LogP values, lower TPSA, and minimal toxicity alerts, making them well-suited for direct topical formulation. Mahanine satisfied multiple physicochemical criteria and its elevated lipophilicity (LogP *>* 6), while requiring formulation consideration, is also consistent with partitioning into lipid-rich skin layers, a property that can be leveraged through lipid-based carrier systems. Annomontine showed acceptable permeability characteristics; its moderately elevated AMES score indicates a parameter to be monitored during formulation development rather than a disqualifying feature at this stage of prioritization. Tricin and tamarixetin, both flavonoids with high TPSA values (*>* 100Å^2^), are recognized candidates for carrier-assisted delivery strategies such as nanoparticle encapsulation or penetration enhancer co-formulation, approaches well established for polar phytochemicals in topical applications. Taraxasterol, with very high lipophilicity (log *P* = 8), is similarly suited to lipid-based nanocarrier systems that can exploit its hydrophobic character for sustained dermal retention. Taken together, these profiles indicate that while protopine and atractylon are directly compatible with passive delivery, the remaining five candidates are well-positioned for formulation-assisted topical application. The variation in physicochemical properties across the panel also highlighted the potential value of combining compounds with complementary delivery profiles, providing the rationale for the pairwise synergy analysis described in the following section.

### 2.6 In silico pairwise synergy prediction using DeepSynergy

All 21 possible two-compound combinations among the seven prioritized phytochemicals were evaluated using a DeepSynergy-inspired framework (Section 3.7) [29]. The complete synergy score matrix is provided in Supplementary File 6, and pairwise scores across all four reference models are visualized in Figure 4.

**Fig. 4:**
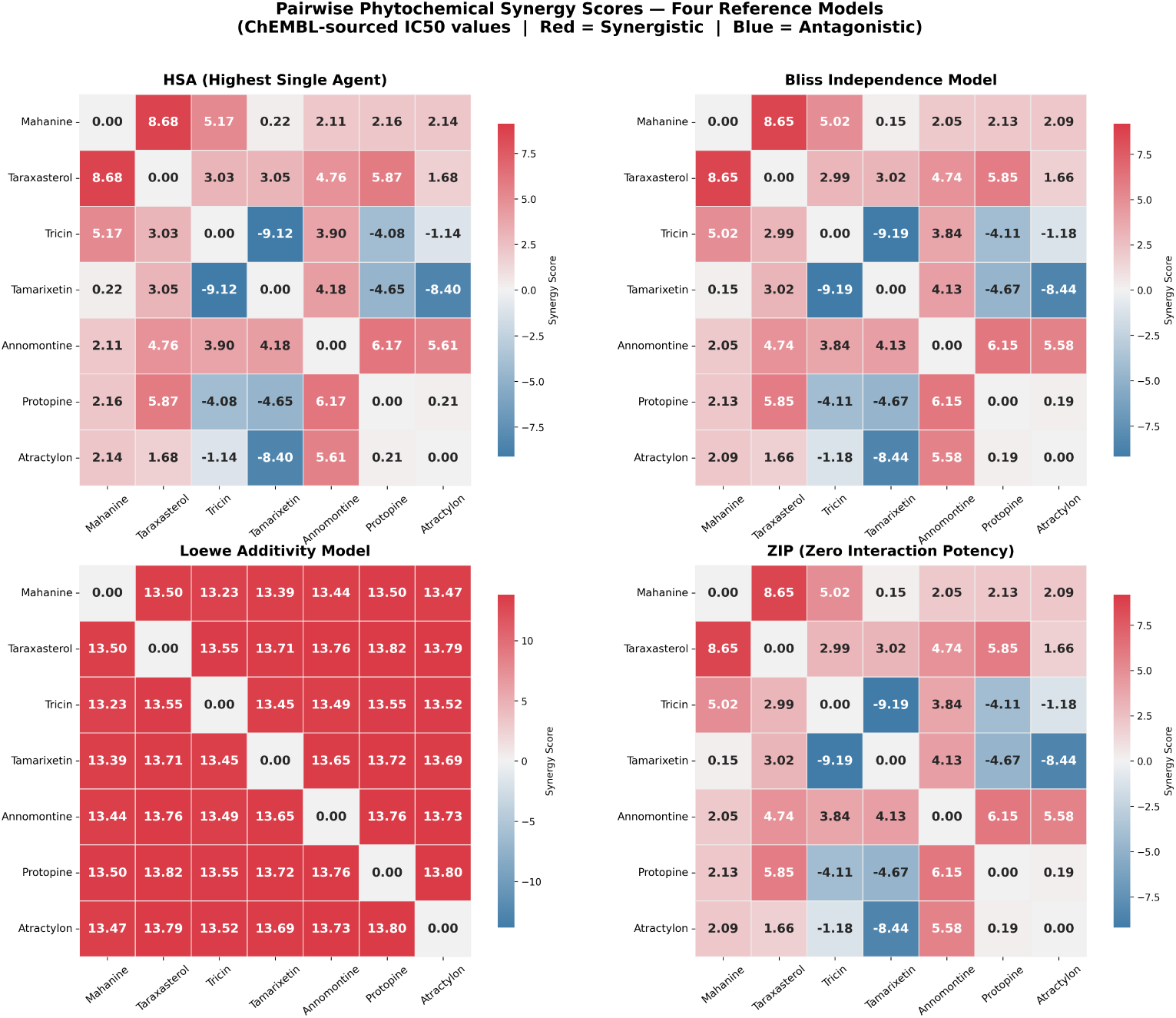
Pairwise phytochemical synergy scores computed under four reference models. Each heatmap represents one scoring model: **(A)** HSA, **(B)** Bliss, **(C)** Loewe, and **(D)** ZIP. Scores are averaged across the full 7 × 7 concentration matrix (0-1000 nM). Red indicates synergy; blue indicates antagonism. Values annotated in each cell represent the mean synergy score for that pair.

Of the 21 pairs evaluated, five exceeded the ZIP synergy threshold of +5, and two pairs fell below −5, indicating predicted antagonism. The highest synergy was observed for Mahanine-Taraxasterol (ZIP = 8.65), followed by Annomontine-Protopine (ZIP = 6.15), Taraxasterol-Protopine (ZIP = 5.85), Annomontine-Atractylon (ZIP = 5.58), and Mahanine-Tricin (ZIP = 5.02). All five synergistic pairs showed cross-model concordance, with HSA and Bliss scores also exceeding +5, representing high-confidence predictions (Table 2). The two predicted antagonistic pairs were Tricin-Tamarixetin (ZIP = −9.19; HSA = −9.12) and Tamarixetin-Atractylon (ZIP = −8.44; HSA = −8.40), both showing strong concordance across HSA and Bliss models. Notably, Annomontine appeared as a synergistic partner in two of the five top-scoring combinations, consistent with its structurally dissimilar profile (Tanimoto ≤ 0.31) relative to its paired compounds, supporting the hypothesis that structural divergence underlies complementary target engagement. Loewe scores were uniformly high across all pairs (≈ 13-14), an artefact arising from the test concentrations (1-1,000 nM) being substantially lower than the compound IC_50_ values (7,000-34,000 nM), which produces a combination index ≪ 1 for all pairs; Loewe scores were therefore excluded from synergy classification and are reported for completeness only.

**Table 2:** Pairwise phytochemical combinations with predicted ZIP synergy score *>* +5 or *<* −5, ranked by ZIP score. All five synergistic pairs showed cross-model concordance (HSA and Bliss both *>* +5). Loewe scores are reported but not used for classification (see text). Tanimoto: pairwise structural similarity coefficient.

| Pair | Tanimoto | HSA | Bliss | Loewe<sup>†</sup> | ZIP |
| --- | --- | --- | --- | --- | --- |
| *Synergistic pairs (ZIP $> +5$)* |  |  |  |  |  |
| Mahanine + Taraxasterol | 0.253 | 8.68 | 8.65 | 13.50 | 8.65 |
| Annomontine + Protopine | 0.311 | 6.17 | 6.15 | 13.76 | 6.15 |
| Taraxasterol + Protopine | 0.272 | 5.87 | 5.85 | 13.83 | 5.85 |
| Annomontine + Atractylon | 0.251 | 5.61 | 5.58 | 13.73 | 5.58 |
| Mahanine + Tricin | 0.455 | 5.17 | 5.02 | 13.23 | 5.02 |
| *Antagonistic pairs (ZIP $< -5$)* |  |  |  |  |  |
| Tricin + Tamarixetin | 0.979 | -9.12 | -9.19 | 13.45 | -9.19 |
| Tamarixetin + Atractylon | 0.620 | -8.40 | -8.44 | 13.69 | -8.44 |

<sup>†</sup>Loewe scores are not used for classification (see text for explanation).

### 2.7 Cross-mapping of IPA pathway signals with phytochemical-targeted genes

#### Canonical pathway enrichment

IPA canonical pathway analysis identified enrichment of immune and barrier-related pathways (see Supplementary File 7), including antimicrobial peptides, interferon *α/β* signaling, interferon *γ* Signaling, keratinization, OAS antiviral response, and role of IL-17A in psoriasis (Figure 5A). Among all the phytochemical gene targets, *LCN2* mapped to the antimicrobial peptides pathway, *EGR1* mapped to interferon *α/β* signaling, and *CXCL8* mapped to the IL-17A pathway. These genes were identified as downstream targets of the upstream regulator *CREB1* in IPA analysis, a target of the prioritized phytochemical atractylon.

**Fig. 5:**
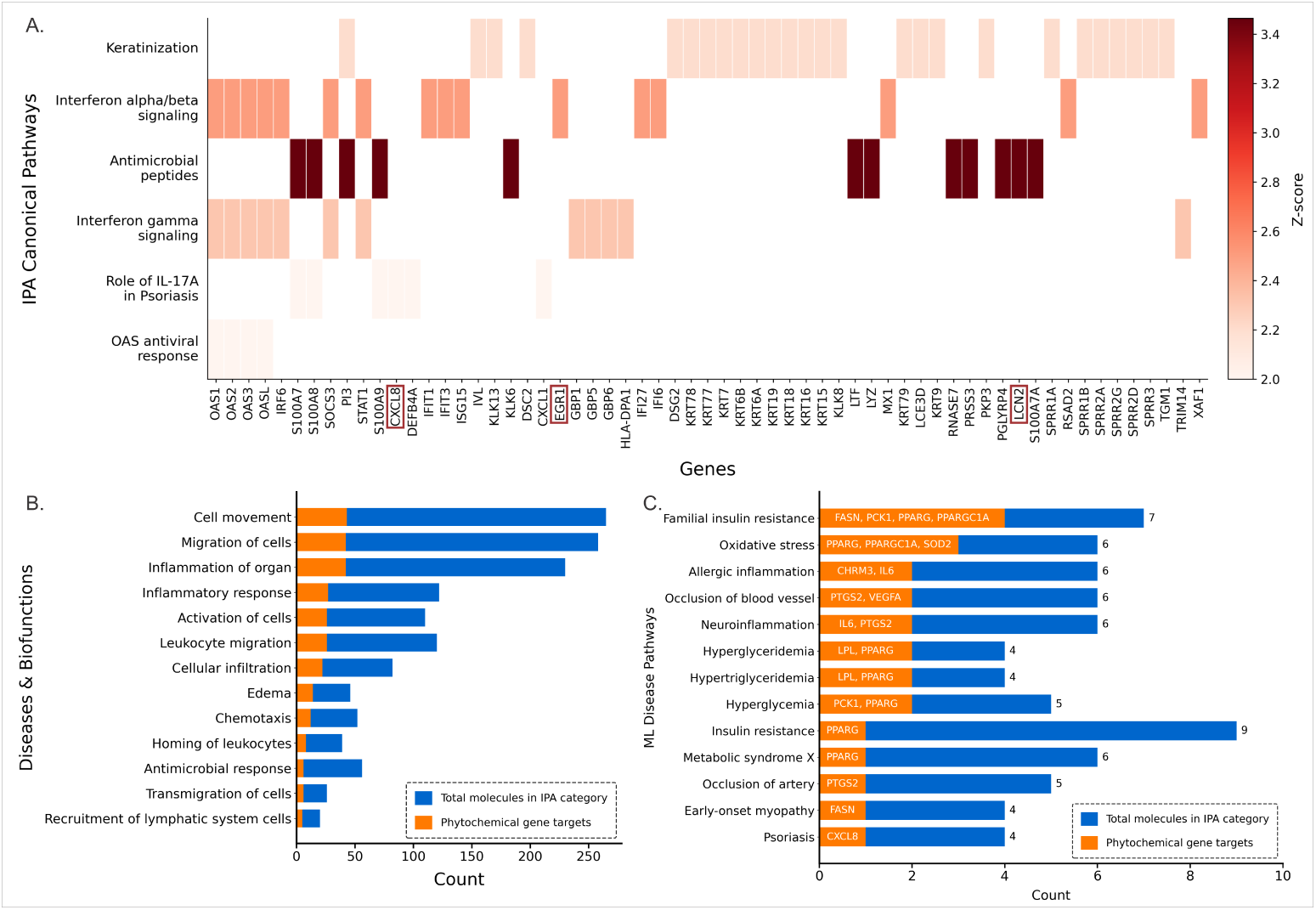
IPA canonical pathway, disease biofunction, and ML disease pathway enrichment of psoriasis DEGs and phytochemical gene targets. **(A)** Heatmap of psoriasis DEGs across enriched IPA canonical pathways (Z-score: 2.0-3.4). Boxed labels indicate phytochemical gene targets (*CXCL8*, *EGR1*, *LCN2*). **(B)** IPA Disease and Biofunction categories showing total pathway genes (blue) and phytochemical gene targets (orange). **(C)** ML disease pathway enrichment showing phytochemical gene overlap (orange) with annotated contributing genes across immunometabolic and comorbidity-associated categories.

#### Diseases & Biofunction and ML disease pathway enrichment

IPA diseases & biofunction analysis groups dataset genes into known biological processes and disease terms, with enrichment confirmed by *p-value* and activation *z-score*. Complementing this, ML Disease pathway analysis uses IPA algorithms trained on large biomedical datasets to identify comorbidity-linked gene associations that fall outside traditional curated pathways. This analysis revealed significant enrichment of immune activation and cell migration-related categories (Figure 5B). For each enriched category, we quantified the number of phytochemical-targeted genes (GoI) relative to the total number of genes in that category. The highest activation *z-scores* for IPA diseases and biofunction were observed for inflammatory response (z = 3.854; 27 GoI of 122 genes), chemotaxis of leukocytes (z = 3.757; 8 of 37), T cell migration (z = 3.701; 5 of 47), migration of mononuclear leukocytes (z = 3.696; 11 of 62), and cell movement of T lymphocytes (z = 3.663; 5 of 44). Broad migration-associated terms also showed strong enrichment, including Cell movement (z = 2.655; 43 of 265), Migration of cells (z = 3.045; 42 of 258), leukocyte migration (z = 3.382; 26 of 120), and inflammation of organ (z = 2.012; 42 of 230). Phytochemical-targeted genes, including *MMP9*, *PTGS2*, and *CYP1A1*, recurrently mapped to inflammatory and migration-related categories. Cell-cycle genes *CCNB1* and *CDK1* were associated with activation and movement-related functions. The complete result is given in Supplementary File 8.

ML disease pathway analysis identified enrichment of immunometabolic and comorbidity-associated categories (see Supplementary File 9). Overlap was quantified based on phytochemical-targeted genes present within each ML-defined pathway. Significant categories included insulin resistance, oxidative stress, neuroinflammation, allergic inflammation, hypertriglyceridemia/hyperglyceridemia, hyperglycemia and vascular occlusion-related terms (Figure 5C). Recurrent overlapping genes across these categories included *PPARG*, *PPARGC1A*, *PCK1*, *FASN*, *IL6*, *PTGS2*, *SOD2*, *LPL*, *VEGFA*, and *CXCL8*. Among the prioritized phytochemicals, atractylon, which targets CREB1, an upstream regulator, overlapped with the insulin resistance module, while tricin and tamarixetin, which target PTGS2, IL6 and VEGFA, overlapped with vascular occlusion categories. This pattern suggests that the phytochemical targets identified here extend beyond cutaneous inflammation into pathways relevant to psoriasis-associated cardiometabolic comorbidities.

### 2.8 PPI Network Construction and Hub Gene Identification in Psoriasis-Associated Targets

To assess interaction patterns, we performed PPI network analysis among selected psoriasis-associated targets (Refer to Methods Section 3.8). Five DEGs (*CCNB1*, *CDK1*, *MMP9*, *PTGS2*, *CYP1A1*) and the upstream regulator *CREB1*, along with their downstream transcriptional targets, were analyzed using STRING (v12.0). The resulting network was imported into cytoscape for visualization and topological analysis. The network showed a highly interconnected structure (Figure 6) where ranking in cytoHubba identified *JUN*, *FOS*, *CREB1*, *IL6*, *CXCL8*, *PPARG*, *MMP9*, and *PTGS2* as top-ranked nodes. Two major regulatory clusters were apparent: an AP-1 transcriptional module formed by *FOS* and *JUN*, and a *CREB1* centered module linking the upstream regulator to its downstream targets.

**Fig. 6:**
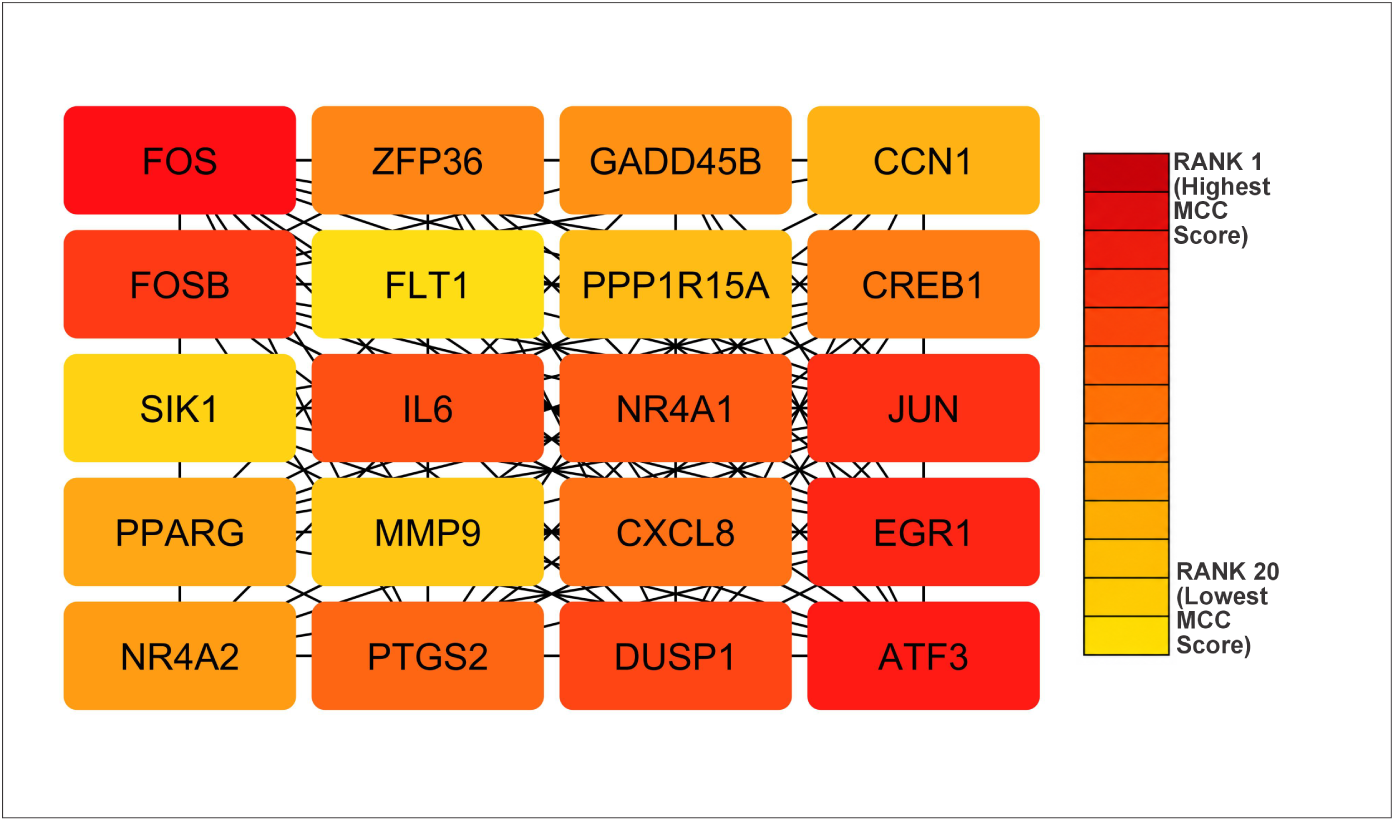
Top-ranked hub genes from PPI network analysis based on MCC scores. Color intensity represents relative centrality, with red indicating higher hub significance.

### 2.9 Tripartite network linking phytochemicals, gene targets, and psoriasis-associated pathways

A tripartite network was constructed by mapping the gene targets of each prioritized phytochemical to psoriasis-associated pathways identified from enrichment analysis (Refer to Methods Section 3.9), retaining only intersecting pathways for inclusion in Figure 7. For complete result, refer to Supplementary File 10.

**Fig. 7:**
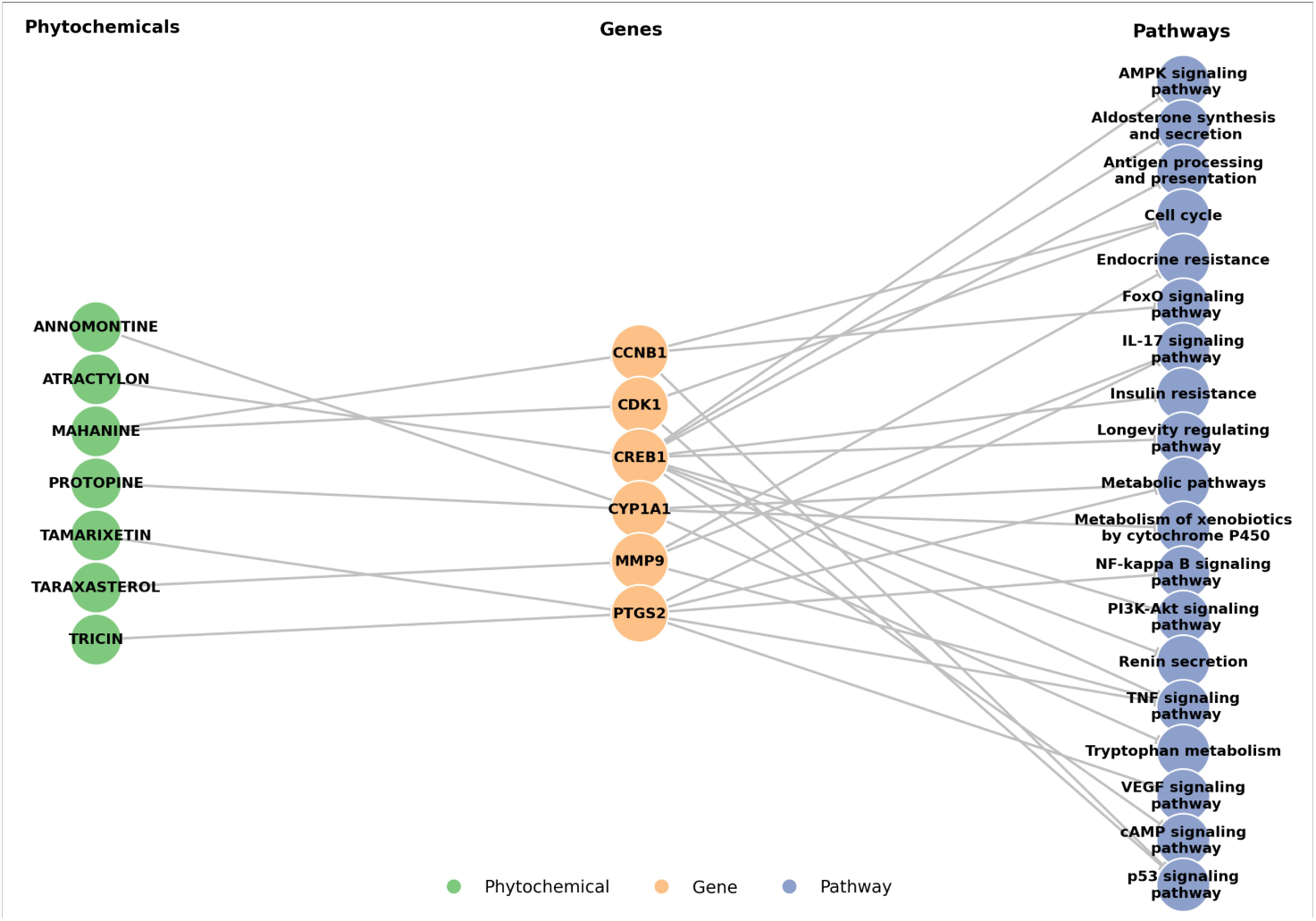
Tripartite network linking phytochemicals, their gene targets, and psoriasis-associated pathways. The network is organized into three layers: phytochemicals (green), gene targets (orange), and pathways (blue). Edges represent reported interactions.

The network includes three layers: (i) prioritized phytochemicals (mahanine, atractylon, protopine, annomontine, taraxasterol, tricin, and tamarixetin); (ii) their gene targets (*CCNB1*, *CDK1*, *CREB1*, *CYP1A1*, *MMP9*, *PTGS2*); and (iii) intersecting psoriasis-associated pathways.

We observed that *Mahanine* targets *CCNB1* and *CDK1*, connecting to cell cycle, FoxO, and p53 signaling pathways. *Atractylon* targets *CREB1*, linking to PI3K-Akt, AMPK, and TNF signaling pathways. *Protopine* and *Annomontine* target *CYP1A1*, mapping to retinol metabolism, tryptophan metabolism, and cytochrome P450-mediated xenobiotic metabolism pathways. *Taraxasterol* targets *MMP9*, connecting to IL-17 and TNF signaling pathways. *Tricin* and *Tamarixetin* target *PTGS2*, mapping to IL-17, NF-*κ*B, and VEGF signaling pathways.

Taken together, the seven prioritized phytochemicals converge on six shared gene targets that collectively intersect with at least ten psoriasis-relevant signaling pathways, a notably broad pathway coverage for a small set of natural compounds.

## 3 Methods

In this study, we applied a structured workflow to identify and prioritize multi-target phytochemicals modulating psoriasis-associated genes. Figure 8 summarizes the overall analytical workflow used in this study.

**Fig. 8:**
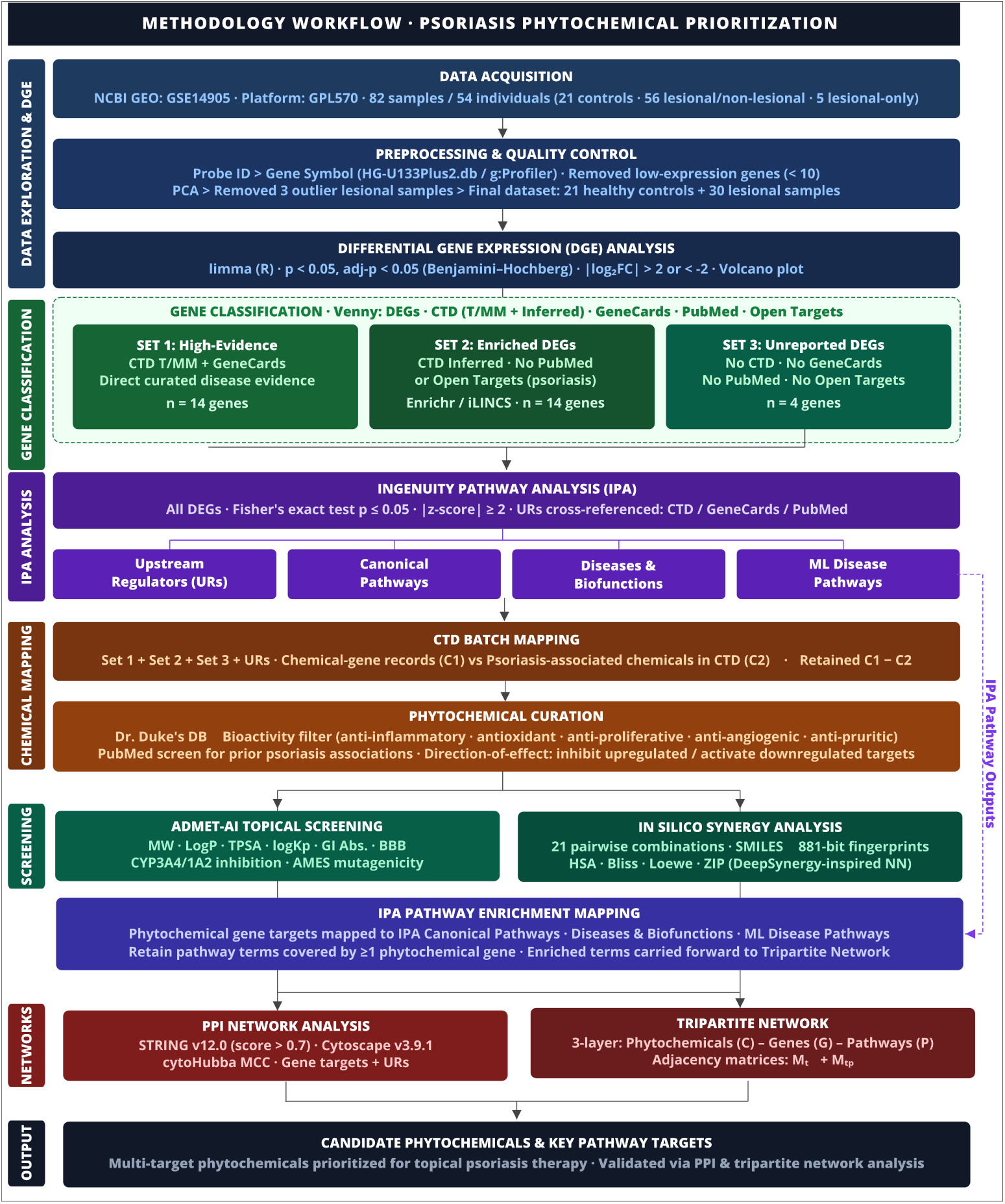
Workflow for the systematic identification and screening of phytochemical candidates. The study began with dataset selection and data preprocessing, followed by DGE analysis. Identified genes were categorized into previously unreported and high-evidence groups. Functional characterization involved pathway and regulator analysis via iLINCS and IPA. Candidate genes were identified through chemical-gene mapping (CTD) with specific phytochemical filtering. Final screening included ADMET-based topical suitability and ZIP-based synergy analysis, concluded by PPI network and enrichment analysis to validate therapeutic targets.

### 3.1 Data Preprocessing and Exploratory Analysis

Gene expression profiles were retrieved from the NCBI GEO repository *GSE14905*, generated using the microarray platform (GPL570) [25]. The dataset comprised 82 samples obtained from 54 individuals, including 21 healthy controls, 28 psoriasis patients with paired lesional and non-lesional biopsies (56 samples), and 5 additional lesional-only samples. Probe IDs were converted to gene symbols using the HG-U133Plus2.db annotation via the g:profiler platform [30]. Further, data were cleaned by removing genes with low expression values (*<* 10) across samples. Then PCA was performed on the normalized gene expression matrix to evaluate sample clustering and detect outliers (Figure 1A). Three lesional samples were identified as outliers and removed for downstream analysis [31]. The final dataset consisted of 21 healthy control samples and 30 lesional samples.

### 3.2 Differential Gene Expression Analysis

Differential Gene Expression Analysis (DGE) analysis was performed using the limma package in R [32]. Genes were considered statistically significant if *p-value* was less than 0.05, *adj p-value* less than 0.05 (calculated using the Benjamini-Hochberg (BH) procedure to control the false discovery rate), and log_2_ fold change greater than 2 or less than -2 [33]. A volcano plot was generated from the resulting statistics to visualize DEG distribution.

### 3.3 Identification of literature-undocumented and high-evidence DEGs

The DEGs were cross-referenced with psoriasis-associated genes in CTD [26] and GeneCards [27]. CTD organizes gene-disease associations into two categories: therapeutic/marker/mechanism (T/MM) and inferred associations. T/MM covers genes with direct curated evidence as disease markers, drug targets, or pathway components, and inferred associations, reflects indirect relationships drawn from chemical-gene-disease interaction records. GeneCards was used as a complementary resource, as it consolidates genomic, transcriptomic, and functional annotation for human genes, including known disease links. Gene overlaps across the three sources were identified using *Venny* [28]. Exclusive DEGs and the ones that overlapped with CTD inferred associations were then screened in PubMed and the Open Targets platform to determine whether any direct psoriasis association existed for each gene. This screening produced three working gene sets:

- Set 1: high-evidence genes - DEGs present in both CTD T/MM and GeneCards.
- Set 2: DEGs overlapping with CTD inferred with no record in PubMed, or Open Targets.
- Set 3: previously unreported genes - DEGs with no record in CTD, GeneCards, PubMed, or Open Targets in the context of psoriasis.

Further, functional enrichment analysis was performed on Set 2.

### 3.4 Ingenuity Pathway Analysis (IPA)

We performed pathway analysis on IPA (Qiagen), which is a web-based platform for analyzing gene expression data using curated biological pathway databases, upstream regulator inference, and disease annotation [34]. DEGs with log_2_ fold-change and adjusted *p <* 0.05 value were imported for pathway and regulatory analysis. IPA generates four analytical outputs - *Upstream Regulators*, *Canonical Pathways*, *Diseases & Biofunctions* and *ML Disease Pathways*. Upstream Regulators (URs) are genes which are upstream in a signaling pathway and can effect the expression of other genes. *Canonical Pathways* are curated biological pathway maps. *Diseases & Biofunctions* links DEGs to biological processes and disease terms, with predicted activation or inhibition state. *ML Disease Pathways* are computationally derived disease associations inferred from large biomedical datasets, capturing comorbidity-linked gene patterns. From IPA those results were retained where enrichment significance value (Fisher’s exact test *p* − *value*) is ≤ 0.05 and predicted activation state (*z-score*) is *z* ≥ +2 and *z* ≤ −2. *z* ≥ +2 indicates activation and *z* ≤ −2 indicates inhibition.

Further, we cross-referenced URs with genes in CTD, GeneCards, and PubMed to identify regulators with previously unreported associations with psoriasis. Later, the genes present in IPA results were compared against the phytochemical gene targets to identify the enriched terms targeted by phytochemicals.

### 3.5 CTD Batch Mapping and Phytochemical Curation

Chemical-gene interaction records were retrieved from CTD using batch queries for all three gene sets - Set 1, Set 2 and Set 3 and URs. Retrieved chemicals (C1) were compared with psoriasis-associated chemicals (C2) in CTD to identify chemicals without prior psoriasis associations (C1-C2) Chemicals were further filtered using Dr. Duke’s Phytochemical and Ethnobotanical Database [35] to retain phytochemicals with documented bioactivities, such as anti-microbial, anti-inflammatory, anti-proliferative, antioxidant, anti-angiogenic, anti-pruritic. Filtered phytochemicals were screened using PubMed to determine previously reported associations with psoriasis. For each phytochemical-gene pair, reported molecular effects (inhibition or activation of the target; increase or decrease in mRNA or protein expression) were compiled. Phytochemicals were prioritized based on direction-of-effect consistency, defined as inhibition of upregulated targets or activation of downregulated targets.

### 3.6 ADMET-AI Topical Suitability Screening

Shortlisted phytochemicals were evaluated for topical suitability using *ADMET-AI* [36] across physicochemical, pharmacokinetic, solubility, and toxicity parameters. The filtering criteria, summarised in Table 3, were adapted from SwissADME guidelines [37], with relaxed thresholds applied for low-percentage formulations (*<*1-2%) [38].

**Table 3:**
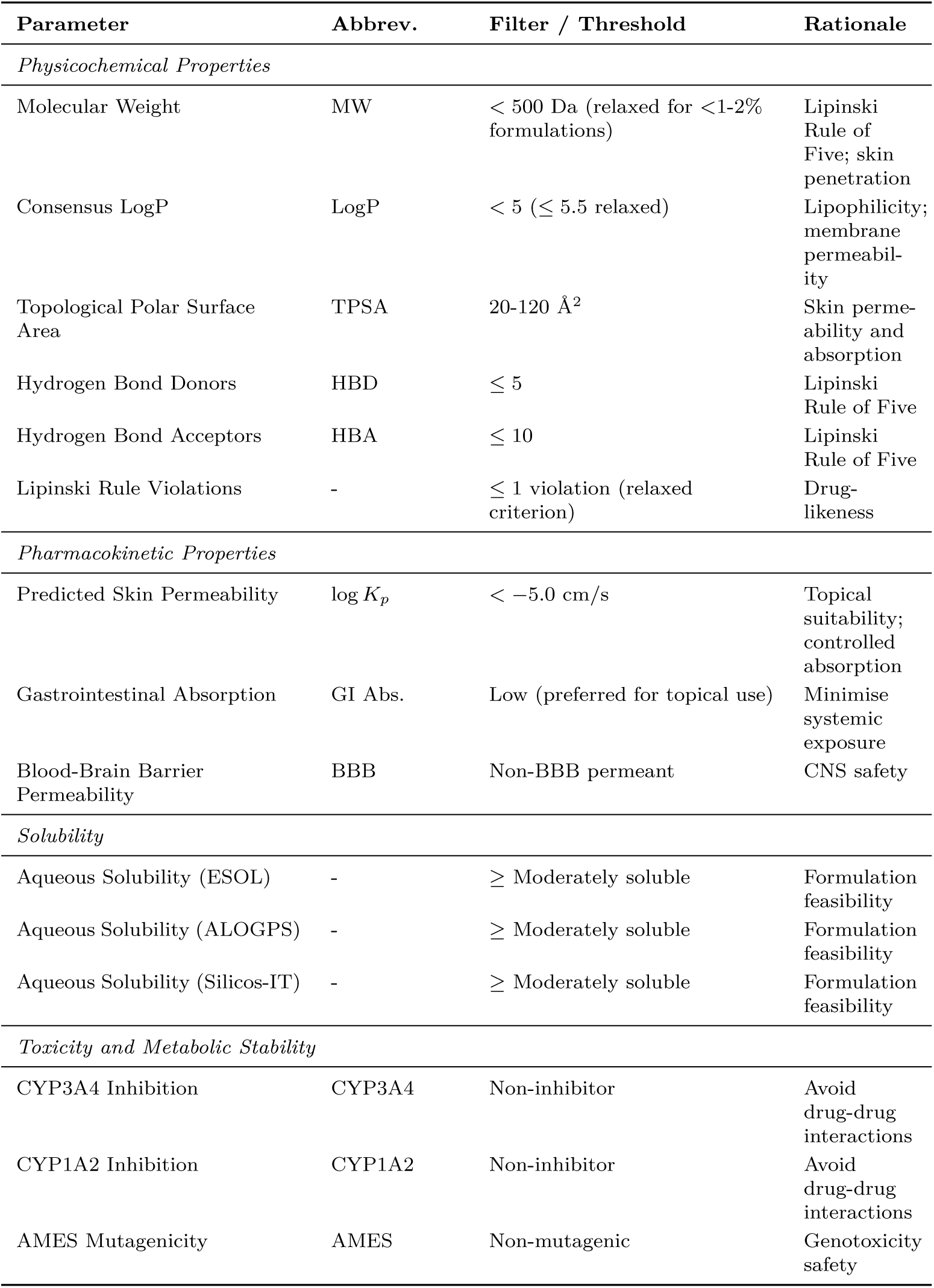
ADMET Screening Parameters and Filter Criteria Adapted from SwissADME Guidelines.

### 3.7 In Silico Synergy Analysis

Pairwise synergy among the seven prioritized phytochemicals was assessed across all 21 possible combinations using a DeepSynergy-inspired framework [29]. Canonical SMILES and 881-bit CACTVS molecular fingerprints were retrieved from Pub-Chem [39], and pairwise structural similarity was quantified using the Tanimoto coefficient. IC_50_ values were sourced from ChEMBL [**?**] for six compounds (Mahanine, 7,000 nM, PMID: 23829449; Tricin, 8,000 nM, PMID: 27955927; Tamarixetin, 14,850 nM, PMID: 25139569; Annomontine, 19,120 nM, PMID: 36931118; Protopine, 34,000 nM, PMID: 20594848; Atractylon, 25,100 nM, PMID: 9544564); Taraxasterol (32,000 nM) required a structural-class estimate as no ChEMBL record was available. Single-agent effects were modelled using the Hill equation (*h* = 1.5) across a 7 × 7 concentration grid (1-1,000 nM). Synergy was quantified using four reference models HSA, Bliss, Loewe, and ZIP [40] with ZIP as the primary criterion (*>*+5: synergistic; *<*−5: antagonistic) [41]. A three-layer feedforward neural network (input: concatenated fingerprint pairs; architecture: 1,762→64→16→1; ReLU activations; 600 epochs) was trained on computed ZIP scores to provide internal consistency validation, with agreement assessed by Pearson *r*. For complete implementation refer to the [https://github.com/rishemjit/Psoriasis]. All results are *in silico* predictions requiring experimental validation in psoriasis-relevant cell-based assays.

### 3.8 Protein-Protein Interaction Network Analysis and Functional Enrichment

) We retrieved interaction data for selected gene targets and upstream regulators from the STRING database (v12.0) using a high-confidence interaction score threshold of > 0.7 [42]. The interaction network was visualized and analyzed using Cytoscape v3.9.1 [43]. Node centrality was calculated using the cytoHubba plugin [44] with the Maximal Clique Centrality (MCC) algorithm, a method that identifies central proteins based on how deeply embedded they are within tightly connected subnetworks.

The protein-protein interaction network was formally defined as a graph

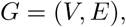

where *V* represented the set of genes (nodes) and *E* represented the set of high-confidence interactions (edges) derived from STRING.

### 3.9 Construction of a Phytochemical-Gene-Pathway Tripartite Network

A tripartite network was constructed to integrate prioritized phytochemicals, their gene targets, and significantly enriched psoriasis-associated pathways. Gene targets were mapped to enriched pathways, and only overlapping associations were retained. The network consisted of three node sets: phytochemicals (*C*), gene targets (*G*), and enriched pathways (*P*). Gene-phytochemical and gene-pathway relationships were represented using adjacency matrices:

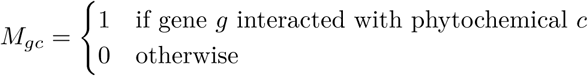

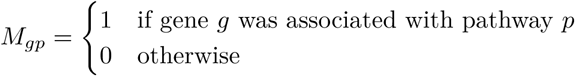

The tripartite graph *T* = (*C, G, P, E*) was assembled from these relationships for network visualisation.

## 4 Discussion

This study applied a computational framework to characterize gene expression changes in psoriatic skin and identify phytochemical candidates for topical intervention. Our analysis highlights coordinated activation of interferon-associated programs alongside known IL-17/TNF inflammatory circuits, consistent with a disease process that extends well beyond skin-confined pathology. Psoriasis operates as a systemic inflammatory disorder in which immune signaling intersects with antiviral responses and metabolic stress pathways [45]. The co-activation of these programs in lesional skin aligns with established roles for the IL-17/TNF axis in keratinocyte activation and leukocyte recruitment [46], and is consistent with transcriptomic profiling of psoriatic plaques reported in prior studies [47]. Type I interferon and nucleic-acid-sensing modules were among the most prominently activated programs in the dataset, with *STAT1*, *IRF1*, *IRF3*, and *IRF7* emerging as central predicted regulators. This pattern is in line with prior reports of interferon-stimulated gene programs and antimicrobial peptide expression in psoriatic plaques [48, 49]. Notably, type III interferons (*IFNL1*, *IFNL4*) were also predicted as active upstream regulators, a finding that distinguishes this analysis from literature [16] which focused primarily on type I interferon axes. Unlike type I interferons, which have well-documented roles in psoriasis, type III interferons signal through the IFNLR1/IL10RB receptor complex expressed predominantly on epithelial cells including keratinocytes [50]. While their general involvement in psoriatic inflammation has been noted [51], their specific role at the psoriatic epithelial-immune interface remains poorly characterized, positioning them as a potentially novel regulatory axis worth investigating. Taken together, these findings suggest that psoriatic lesions reflect convergent immune programs across multiple pathway axes rather than a single dominant mechanism [52].

Our analysis captured metabolic and oxidative stress signatures in lesional skin that are consistent with the systemic features of psoriasis. ML-derived disease pathways identified through IPA highlighted oxidative stress, insulin resistance, and lipid metabolism terms, findings that align with the well-established epidemiological links between psoriasis and cardiometabolic comorbidities [53]. Chronic inflammatory signalling is known to promote oxidative imbalance and mitochondrial stress with down-stream systemic effects [54]. These patterns collectively support an immunometabolic framework in which interferon programs intersect with IL-17/TNF signalling and systemic metabolic regulation, and underscore the multi-module nature of psoriatic pathobiology.

At the network level, PPI analysis of our DEG set condensed the transcriptional signal into a hub-centered architecture linking inflammatory, angiogenic, and metabolic pathways within a compact regulatory core. Hub-dominated network organisation has been described in other chronic inflammatory settings and may reflect self-reinforcing regulatory circuits that sustain pathway activity over time [55]. In our analysis, AP-1 transcription factors (*JUN* and *FOS*) and *CREB1* emerged as the most central transcriptional integrators. Prior work supports roles for AP-1 in keratinocyte differentiation and IL-23-linked chemokine programs [56], and for *CREB1* in stress response and inflammatory regulation across immune cell types [57]. The enrichment results obtained here position these hubs at the intersection of MAPK, NF-*κ*B, and cytokine signaling, connecting inflammatory stimuli to proliferation, angiogenesis, oxidative stress, and lipid homeostasis [58]. Additional hub nodes *MMP9* and angiogenic mediators (*VEGFA*, *FLT1*) identified in this network are consistent with dermal remodeling and vascular expansion observed in lesional skin [59], while *PPARG* and *PTGS2* connect lipid and eicosanoid metabolism with cytokine amplification and keratinocyte proliferation [60].

The seven prioritized phytochemicals i.e., mahanine, taraxasterol, tricin, tamarixetin, annomontine, protopine, and atractylon, mapped to multiple regulatory nodes across inflammatory, proliferative, and metabolic layers of this network. In the tripartite network constructed here, mahanine connected to cell-cycle regulators (*CCNB1*, *CDK1*); taraxasterol to matrix remodeling through *MMP9* ; tricin and tamarixetin to *PTGS2* within NF-*κ*B-linked inflammatory and angiogenic modules; atractylon to *CREB1* and PI3K-Akt/AMPK pathways; and protopine and annomontine to *CYP1A1* within xenobiotic and retinoid metabolism modules. These associations are derived from curated interaction databases and define mechanistic hypotheses for experimental testing rather than confirmed pharmacological activities.

For topical development, ADMET-AI screening confirmed that all seven prioritized phytochemicals carry profiles supporting further development, with protopine and atractylon demonstrating physicochemical properties most directly compatible with passive dermal penetration [61]. For the remaining candidates, the physicochemical characteristics identified high lipophilicity in mahanine and taraxasterol, and elevated polarity in the flavonoids tricin and tamarixetin are well-recognized as addressable through established formulation strategies, including lipid-based nanocarriers, penetration enhancers, and encapsulation systems, rather than as barriers to development [61]. The variation in delivery profiles across the panel further motivated exploration of compound combinations, since pairing agents with complementary physicochemical and mechanistic properties can improve overall pathway coverage and skin exposure. The *in silico* synergy analysis confirmed that pairwise combinations within this phytochemical panel are broadly compatible: no pair was predicted to be antagonistic, and eight combinations exceeded the synergy threshold. The recurring involvement of atractylon across the top-ranked pairs is consistent with its multi-target profile, suggesting mechanistic complementarity with both alkaloid and flavonoid partners. Taken together, the ADMET and synergy data provide converging computational support for a combination-based topical strategy, with experimental confirmation in psoriasis-relevant cell-based assays as the next required step.

## 5 Conclusion

This study demonstrates that psoriasis is driven by a convergent multi-module regulatory network encompassing type I/III interferon signaling, IL-17/TNF inflammatory circuits, and immunometabolic dysregulation rather than a purely cytokine-restricted process. Integrating transcriptomic analysis, network modeling, and phytochemical mapping revealed regulatory hubs that connect inflammatory and metabolic pathways within psoriatic lesions. Using this framework, several natural compounds were prioritized as potential multi-target modulators with relevance for topical therapeutic strategies. Together, these findings demonstrate how systems-level analysis can link disease transcriptomics with mechanism-guided phytochemical prioritization. Future experimental studies in keratinocyte and skin models will be necessary to determine whether these compounds can effectively modulate psoriasis-associated signaling.

### 5.1 Data and Code Availability

GitHub repository contains all analysis scripts, processed datasets, and supplementary files generated in this research work: https://github.com/rishemjit/Psoriasis.

## Supplementary information

- **Supplementary File 1:** S_DEGs.xlsx - Complete list of DEGs, including log_2_ fold-changes and adjusted *p*-values used in all downstream analysis.
- **Supplementary File 2:** S_KEGG.xlsx - GO (biological process, molecular function, cellular component), KEGG, and Reactome enrichment results for the 14 candidate genes, semantically reduced using rrvgo. Includes all enriched terms, raw *p-values*, odds ratios, and rrvgo-assigned parent terms.
- **Supplementary File 3:** S_UR.xlsx - IPA upstream regulators (URs) for the DEG set, with activation *z*-scores, overlap *p*-values, and literature-evidence tiers.
- **Supplementary File 4:** S_NG-KPC_ixns.xlsx - Known phytochemicals targeting *novel* psoriasis-associated genes.
- **Supplementary File 5:** S_ADMET_AI.xlsx - ADMET-AI predicted physicochemical and pharmacokinetic properties for shortlisted phytochemicals.
- **Supplementary File 6:** S_Synergy_scores.xlsx - SynergyFinder outputs for all pairwise phytochemical combinations: concentration matrices for Relative Inhibition and ZIP synergy scores with corresponding dose units.
- **Supplementary File 7:** S_CP.xlsx - IPA Canonical Pathways significantly associated with the psoriasis transcriptomic signature.
- **Supplementary File 8:** S_DB.xlsx - IPA Diseases & Biofunctions enriched in the dataset, with activation-state predictions and overlapping genes relevant to psoriatic inflammation and tissue remodeling.
- **Supplementary File 9:** S_ML.xlsx - IPA Machine-Learning (ML) disease pathways linked to psoriasis and comorbid traits, with contributing DEGs.
- **Supplementary File 10:** Tripartite_Network.xlsx - Tripartite network node/edge tables linking shortlisted phytochemicals, their gene targets, and psoriasis-relevant pathways (CTD and curated literature).

## Acknowledgements

We gratefully acknowledge the support by Division of Biotechnology and Molecular Medicine, School of Veterinary Medicine, Louisiana State University.

## Appendix A Functional enrichment of DEGs with inferred CTD associations and no prior psoriasis reporting

Of the 556 DEGs identified, those overlapping with CTD inferred associations were screened individually through PubMed (“<`GENE_SYMBOL`>” AND psoriasis in [All Fields]) and the Open Targets platform [62] to check for any direct prior reporting in psoriasis. Open Targets integrates genetic, transcriptomic, and clinical evidence to score gene-disease associations. From the 22 exclusive DEGs, 4 genes - *DANT2* (−2.562), *GK4P* (+2.618), *PRSS3P1* (+2.224), and *LINC01140* (−2.289) – showed no prior reported association with psoriasis in any of these sources. From the 517 DEGs overlapping with CTD inferred associations, screening through PubMed and Open Targets produced a refined set of 32 genes with no direct psoriasis evidence.

Functional enrichment of these 32 genes was carried out using Enrichr and iLINCS across Gene Ontology (biological process, molecular function, and cellular component), KEGG, and Reactome databases. Significance was assessed using Fisher’s exact test (*p <* 0.05), and results were ranked by combined score. To reduce redundancy among GO terms, semantic clustering was performed using the *rrvgo* Bioconductor package (similarity threshold = 0.7, Wang method), retaining only representative non-overlapping terms [63].

Of the 32 genes, 14 produced at least one statistically significant enrichment term at raw *p <* 0.05 after semantic reduction: *SRD5A3*, *PCGF5*, *IRF6*, *DDX17*, *OSBPL8*, *IMMP2L*, *ADAMTS15*, *KLHL18*, *WDR33*, *ZC3H11A*, *TIAM2*, *MRAS*, *KREMEN1*, and *CA4*. The enriched terms fell into six biologically coherent themes with relevance to psoriatic pathology: N-linked glycoprotein biosynthesis via the dolichol pathway (*SRD5A3*); Polycomb-mediated epigenetic gene silencing (*PCGF5*); post-transcriptional mRNA regulation (*DDX17*, *WDR33*, *ZC3H11A*); lipid and membrane homeostasis (*OSBPL8*); mitochondrial protein processing (*IMMP2L*); and cell cycle regulation via the ubiquitin-proteasome system (*KLHL18*). Additional enrichments for interferon signaling (*IRF6*), RAS/MAPK signaling (*TIAM2*, *MRAS*), and WNT pathway antagonism (*KREMEN1*) were consistent with established psoriasis biology. Since each enriched term is driven by a single gene, these results represent gene-level functional annotations and should not be interpreted as evidence of coordinated pathway activity. All enrichment terms, adjusted *p*-values, and odds ratios are provided in Supplementary File 2. The statistically significant terms for all 14 genes are listed in Table A1.

**Table A1:** Statistically significant enrichment results for the 14 candidate genes (*p <* 0.05, raw). Gene direction: ↑ upregulated; ↓ downregulated in psoriatic lesional skin. **N** = new gene-level association in psoriasis (reference supports gene function); **K** = known psoriasis pathway with a new gene-level association (reference supports pathway involvement in psoriasis).

| Gene | Source | Enriched Term | $p$ | Rel. | Ref. |
| --- | --- | --- | --- | --- | --- |
| *SRD5A3* ↓ | GO BP | Polyprenol Metabolic Process (GO:0016093) | 0.009 | N | [64] |
|  | GO BP | Dolichol-Linked Oligosaccharide Biosynthesis (GO:0006488) | 0.035 | N | [64, 65] |
|  | GO BP | Protein N-linked Glycosylation via Asn (GO:0018279) | 0.044 | N | [64] |
|  | GO BP | Steroid Hormone Biosynthetic Process (GO:0120178) | 0.039 | K | [64] |
|  | GO BP | Androgen Metabolic Process (GO:0008209) | 0.039 | K | [64] |
|  | GO MF | Oxidoreductase Activity, CH-CH Group (GO:0016628) | 0.034 | N | [64] |
|  | Reactome | Defective SRD5A3 causes SRD5A3-CDG | 0.007 | N | [64] |
| *PCGF5* ↑ | GO CC | PRC1 Complex (GO:0035102) | 0.030 | N | [66] |
| *IRF6* ↑ | Reactome | Interferon $\alpha/\beta$ Signalling | 0.037 | K | [67, 68, 16] |
| *DDX17* ↑ | GO BP | miRNA Metabolic Process (GO:0010586) | 0.025 | N | [69] |
|  | GO BP | Alternative mRNA Splicing, via Spliceosome (GO:0000380) | 0.035 | N | [69] |
|  | GO BP | Androgen Receptor Signalling Pathway (GO:0030521) | 0.032 | K | [69] |
| *OSBPL8* ↓ | GO BP | Phosphatidylserine Acyl-Chain Remodelling (GO:0036150) | 0.014 | N | [70] |
|  | GO BP | Acylglycerol Homeostasis (GO:0055090) | 0.046 | N | [70] |
|  | GO BP | Organophosphate Ester Transport (GO:0015748) | 0.044 | N | [70] |
|  | GO CC | Cortical Endoplasmic Reticulum (GO:0032541) | 0.013 | N | [70] |
|  | GO MF | Phosphatidylinositol-4-Phosphate Binding (GO:0070273) | 0.048 | N | [70, 71] |
| *IMMP2L* ↓ | GO BP | Mitochondrial Protein Processing (GO:0034982) | 0.018 | N | [72] |

Table A1 - *continued*
| Gene | Source | Enriched Term | *p* | Rel. | Ref. |
| --- | --- | --- | --- | --- | --- |
|  | GO BP | Signal Peptide Processing (GO:0006465) | 0.021 | N | [72] |
|  | GO CC | Serine-Type Endopeptidase Complex (GO:1905370) | 0.021 | N | [72] |
| *KLHL18* ↑ | GO BP | Positive Regulation of Cell Cycle Phase Transition (GO:1901989) | 0.034 | N | [73] |
|  | GO BP | Positive Regulation of Mitotic Cell Cycle (GO:0045931) | 0.049 | N | [73] |
|  | GO CC | CUL3-RING Ubiquitin Ligase Complex (GO:0031463) | 0.031 | N | [73] |
| *WDR33* ↑ | GO CC | mRNA Cleavage and Polyadenylation Specificity Factor Complex (GO:0005847) | 0.039 | N | [74] |
|  | Reactome | Transport of Mature Transcript to Cytoplasm | 0.019 | N | [74] |
|  | Reactome | Processing of Intronless Pre-mRNAs | 0.048 | N | [74] |
| *ZC3H11A* ↑ | GO BP | poly(A)<sup>+</sup> mRNA Export From Nucleus (GO:0016973) | 0.041 | N | [75] |
|  | Reactome | Transport of Mature Transcript to Cytoplasm | 0.019 | N | [75] |
| *TIAM2* ↑ | GO BP | Small GTPase-mediated Signal Transduction (GO:0007264) | 0.045 | K | [76, 77] |
| *MRAS* ↑ | GO BP | Small GTPase-mediated Signal Transduction (GO:0007264) | 0.045 | K | [76, 77] |
|  | Reactome | Gain-of-function MRAS Complexes Activate RAF Signalling | 0.026 | K | [76] |
| *KREMEN1* ↑ | Reactome | Signalling by LRP5 Mutants | 0.014 | K | [78] |
|  | Reactome | Negative Regulation of TCF-dependent Signalling by WNT Ligand Antagonists | 0.035 | K | [78] |
| *CA4* ↓ | GO MF | Carbonate Dehydratase Activity (GO:0004089) | 0.021 | N | [79] |
|  | KEGG | Nitrogen Metabolism | 0.027 | N | [79] |
|  | Reactome | Reversible Hydration of Carbon Dioxide | 0.039 | N | [79] |

